# A chromosome-scale genome assembly of a *Bacillus thuringiensis* Cry1Ac insecticidal protein resistant strain of *Helicoverpa zea*

**DOI:** 10.1101/2022.04.12.488070

**Authors:** Amanda R. Stahlke, Jennifer Chang, Luke R. Tembrock, Sheina B. Sim, Sivanandan Chudalayandi, Scott M. Geib, Brian E. Scheffler, Omaththage P. Perera, Todd M. Gilligan, Anna K. Childers, Kevin J. Hackett, Brad S. Coates

## Abstract

*Helicoverpa zea* (Lepidoptera: Noctuidae) is an insect pest of major cultivated crops in North and South America. The species has adapted to different host plants and developed resistance to several insecticidal agents, including *Bacillus thuringiensis* (Bt) insecticidal proteins in transgenic cotton and maize. *H. zea* populations persist year-round in tropical and subtropical regions, but seasonal migrations into temperate zones increase the geographic range of associated crop damage. To better understand the genetic basis of these physiological and ecological characteristics, we generated a high-quality chromosome-level assembly for a single *H. zea* male from Bt resistant strain, HzStark_Cry1AcR. Hi-C data were used to scaffold an initial 375.2 Mb contig assembly into 30 autosomes and the Z sex chromosome (scaffold N50 = 12.8 Mb and L50 = 14). The scaffolded assembly was error-corrected with a novel pipeline, polishCLR. The mitochondrial genome was assembled through an improved pipeline and annotated. Assessment of this genome assembly indicated 98.8% of the Lepidopteran Benchmark Universal Single-Copy Ortholog set were complete (98.5% as complete single-copy). Repetitive elements comprised approximately 29.5% of the assembly with the plurality (11.2%) classified as retroelements. This chromosome-scale reference assembly for *H. zea*, ilHelZeax1.1, will facilitate future research to evaluate and enhance sustainable crop production practices.

**Significance:** We established a chromosome-level reference assembly for *Helicoverpa zea*, an insect pest of multiple cultivated crops in the Americas. This assembly of a *Bacillus thuringiensis* insecticidal protein resistant strain, HzStark_Cry1AcR, will facilitate future research in areas such as population genomics and adaptations to agricultural control practices.

## Introduction

*Helicoverpa zea* (Boddie), (Lepidoptera: Noctuidae) is a widespread insect in North and South America (Djaman et al. 2019; Huseth et al. 2021; **fig. 1A**). This pest causes extensive damage to plants including maize, cotton, soybean, and vegetable crops, and feeds on many weedy plants (Degrande and Omoto 2013; Cunningham and Zalucki 2014; Leite et al. 2014; Reay-Jones 2019). Relatively rapid development and short generation times (Fitt 1989), annual long-distance migrations (Reay-Jones 2019; Perera et al. 2020), highly polyphagous larvae that tolerate a range of plant secondary defensive metabolites (Niu et al. 2008; Li et al. 2020), and entry into larval diapause during adverse environmental conditions (Roach and Adkisson 1970; Reynolds et al. 2019) contribute to the severity of *H. zea* as a pest. Furthermore, resistance of *H. zea* to chemical insecticides (McCaffrey 1988; Hamadain and Chambers 2001; Jacobson et al. 2009) and *Bacillus thuringiensis* (Bt) crystalline (Cry) and vegetative insecticidal proteins (Vip) expressed in transgenic cotton (Luttrell and Jackson 2012; Rabelo et al. 2020) and maize (Reisig and Reay-Jones 2015; Dively et al. 2016; Bilbo et al. 2019; Yang et al 2019) has led to difficulties controlling crop damage. Resistance to Bt insecticidal proteins is associated with changes in midgut receptors or signal transduction pathways (Soberón et al. 2009). Evidence suggests *H. zea* Bt Cry1Ac resistance is influenced by point mutations in tetraspanin (Jin et al. 2019) or kinesin (Benowitz et al. 2021) genes, or at several loci (Taylor et al. 2021). These reductions in *H. zea* insecticide susceptibility (Yang et al. 2020; Santiago González et al. 2021) contribute to increased levels of crop damage and lower yields (Reisig and Kurtz 2018; Reay-Jones 2019), presenting a threat to global food security (Coates et al. 2015). This has been exacerbated by the introduction and spread of the sister species, *H. armigera*, into South America and the Caribbean (Czepak et al. 2013; Tay et al. 2013; Arnemann et al. 2015; Murúa et al. 2016; Sosa-Gómez et al. 2016; Tembrock et al. 2019) where introgression of adaptive alleles has led to novel phenotypes that further complicate pest management efforts (Anderson et al. 2018; Cordeiro et al. 2020; Valencia-Montoya et al. 2020; Rios et al. 2021).

**Fig. 1.**
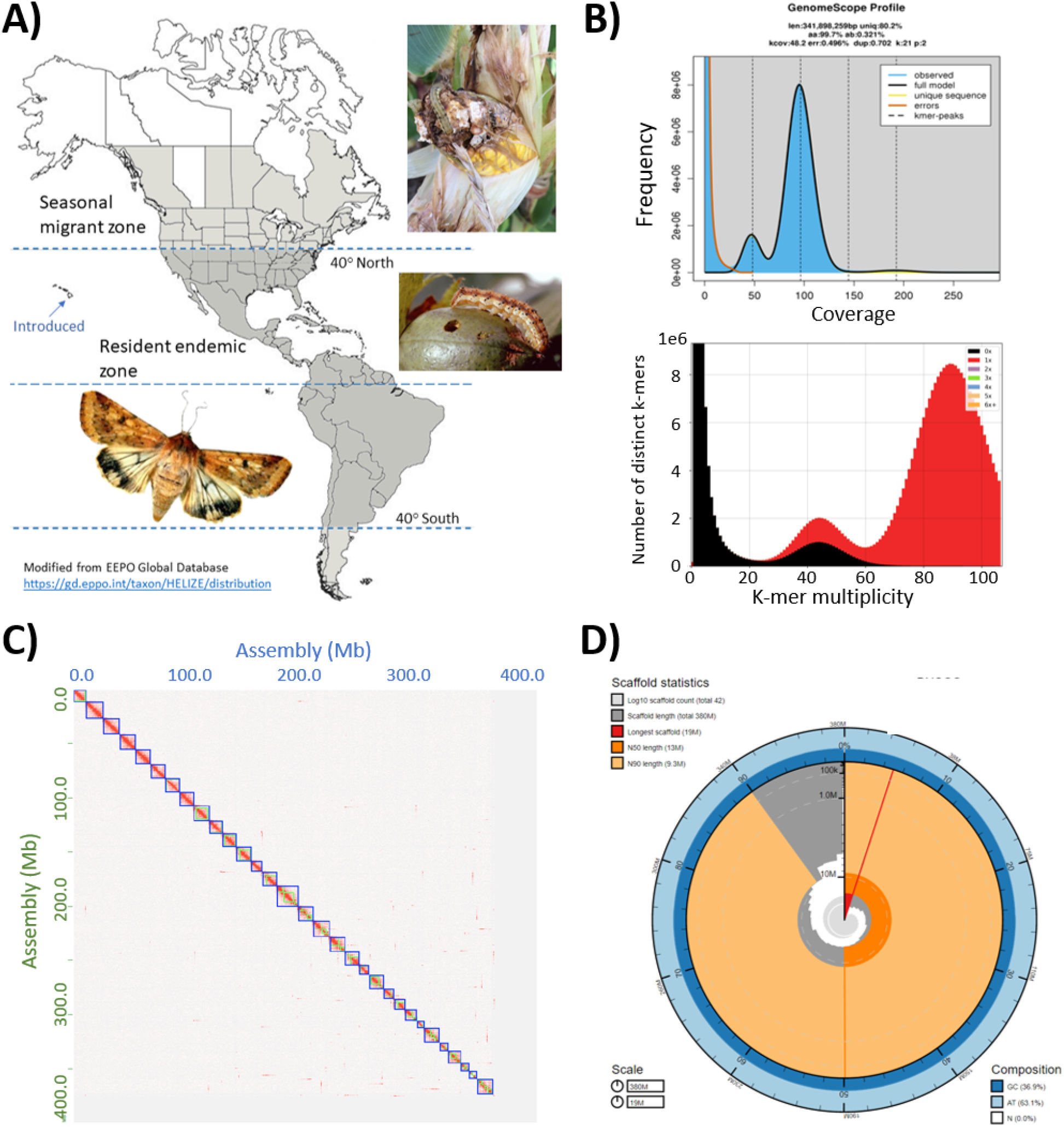
Features of *Helicoverpa zea* biology and genome assembly. *(A)* Approximate geographic distribution and examples of crop damage, *(B)* K-mer analyses of short reads for estimated genome size, heterozygosity, and duplication level. *(C)* Hi-C contact map adjoining and ordering 31 pseudochromosomes. *(D)* Snailplot indicating metrics of 31 pseudochromomes within the entire assembly of 42 scaffolds.

Prior *Helicoverpa* genome sequencing using short-read data produced a 341.1 Mb *H. zea* assembly, Hzea_1.0, arranged in 2,975 scaffolds, and a similar low contiguity 337.1 Mb assembly for the closely related *H. armigera*, Harm_1.0 (Pierce et al. 2017). Although these assemblies have been useful for population analyses (Valencia-Montoya et al. 2020; Taylor et al. 2021), chromosome-scale assemblies empower multiple areas of genomic research (Lewin et al. 2018; Childers et al. 2021). As such, we assembled and scaffolded a high-quality chromosome-scale genome for *H. zea*. Our approach implemented best practices for non-model genome assembly with noisy long-reads, whereby an initial assembly from a single individual produced haplotype-specific contigs, followed by purging of duplicate haplotypes prior to Hi-C scaffolding and manual curation, error correction (polishing) including a mitochondrial assembly, and filtering of possible contaminants (Rhie et al. 2021, Howe et al. 2021). This assembly is substantially more contiguous and complete compared to the prior *Helicoverpa* genome resources and serves as an exemplar for developing high quality resources to improve understanding of insecticide resistance, population dynamics, and efficacy of pest management strategies.

## Results and Discussion

### Genome assembly, scaffolding, annotation, and completeness

In total, 54.0 Gb of raw PacBio continuous long read (CLR) data was generated from a single *H. zea* HzStark_Cry1AcR strain male (supplementary table S1), which resulted in an estimated 148.8-fold coverage based on a 362.8±8.8 Mb flow cytometry estimated genome size (Coates et al. 2017). GenomeScope k-mer analyses of 154.6 million whole-genome shotgun Illumina reads provided an estimated genome size of 341.9 Mb, 80.2% unique (non-repetitive) content, and low heterozygosity (0.3%) (fig. 1B), the latter likely a result of using a single individual from a colony with a degree of inbreeding. Assembly of CLR reads using FALCON-Unzip produced an initial 375.2 Mb assembly consisting of 134 primary pseudo-haploid contigs and 99.4 Mb in 765 alternate contigs (supplementary table S2). After purging duplicate haplotigs, the assembly was improved to 81 primary and 801 alternate contigs (Table 1 and supplementary table S2). These 81 deduplicated primary contigs contained 4.8% fewer duplicated (D) orthologs compared to the initial FALCON-Unzip assembly when assessed with BUSCO.

**Table 1.** *Helicoverpa zea* genome assembly and annotation metrics

| Metric | Value |
| --- | --- |
| Alternate contig assembly |  |
| Total number | 801 |
| Assembly size (bp) | 133,873,018 |
| L50/N50 (bp) | 90/464,725 |
| L90/N90 (bp) | 417/49,392 |
| Largest contig (bp) | 1,812,199 |
| Primary assembly (ilHelZeax1.1)<sup>1</sup> |  |
| Scaffolds |  |
| Total number | 42 |
| Size (bp) | 375,165,593 |
| L50/N50 (bp) | 14/12,841,304 |
| L90/N90 (bp) | 27/9,278,413 |
| Largest (bp) | 18,805,280 |
| Contigs |  |
| Total number | 65 |
| Size (bp) | 375,163,385 |
| L50/N50 (bp) | 15/10,871,041 |
| L90/N90 (bp) | 34/3,915,976 |
| Largest (bp) | 15,512,169 |
| BUSCOs |  |
| Complete (C) | 5227 (98.8%) |
| Complete single copy (S) | 5209 (98.5%) |
| Complete duplicated (D) | 18 ( 0.3%) |
| Fragmented (F) | 17 ( 0.3%) |
| Missing (M) | 42 ( 0.9%) |
| Gene structural annotations |  |
| Genes (total) | 16.988 |
| Protein coding | 14.933 |
| Non-coding | 1,812 |
| Pseudogenes | 242 |
| RefSeq mRNAs | 23,683 |
| RefSeq proteins | 23,696 |
| Masked repeats bp (%)<sup>2</sup> | 110,825,260 (29.5%) |
| Repeat content (%)<sup>3</sup> | 23.6% |

1. *Helicoverpa zea* WGS Project: GenBank accession: JAKYJW000000000
2. See RepeatMasker results in supplementary table S3

Scaffolding of primary contigs using Hi-C data broke five mis-joins in three contigs resulting in 42 total scaffolds (fig. 1C) and improved overall contiguity (scaffold N50 of 12.8 Mbp in 14 scaffolds) in the final ilHelZeax1.1 assembly (fig. 1D, table 1 and supplementary table S2). Hi-C contact mapping between contigs resolved 31 scaffolds, representing the expected set of 30 autosomes and the Z chromosome based on a *H. armigera* karyotype (Sahara et al. 2013) and other noctuid assembly data (supplementary table S3). The remaining 11 scaffolds comprised 1.2 Mb and represented < 0.3% of the entire assembly. Error-correcting the scaffolds with Illumina data using the polishCLR pipeline (Stahlke et al. 2022) increased the consensus quality value (QV) from 38.9 to 42.0, and improved the contig assembly L50 and N50 to 15 and 10.9 Mb, respectively. Scaffold length distributions did not change through polishing. The final scaffolded length of 375.2 Mbp is similar to prior flow cytometry estimates (Coates et al. 2017), and larger and more contiguous than the prior 341.1 Mb Hzea_1.0 assembly (Pearce et al. 2017) or other *Helicoverpa* assemblies (supplementary table S2).

Overall, the ilHelZeax1.1 assembly exceeds the minimum reference standard of 6.C.Q40 (>1.0 Mb contig and >10.0 Mb scaffold N50) set for eukaryotic species by the Earth BioGenome Project (Lewin et al. 2018) and meets the qualifications for chromosome level (7.C.Q50; Lawniczak et al. 2022). Assessing the assembly relative to short-reads, k-mer spectra indicated a moderate degree of heterozygosity in the primary pseudo-haploid assembly, shown by a histogram peak (x = 90) corresponding to both haplotypes that is much greater compared to that for single-copy k-mers (x = 45; fig. 1B). Also, almost no redundant sequence was apparent. Corresponding BUSCOs indicated a high level of representation and minimal duplication (score of C:98.8% [S:98.5%, D:0.3%], F:0.3%, and M:0.9%; table 1), which is similar to chromosome-scale assemblies from other species in the Family Noctuidae (supplementary table S3). The NCBI Eukaryotic Genomic Annotation Pipeline predicted 16,988 genes, of which 14,922 protein coding genes gave rise to 23,683 RefSeq transcripts (table 1). Greater than 94% RefSeq models were supported by RNA-seq evidence. A total of 110.8 Mb of repeats and transposable elements were masked in the ilHelZeax1.1 assembly (supplementary table S4).

### Mitochondrial genome assembly

The mitoPolishCLR pipeline (Formenti and Stahlke 2022) provided a robust assembly and annotation of the *H. zea* mitochondrial genome. The final HzStark_Cry1AcR mitochondria assembly was 15,351 bp and slightly larger than the 15,343 bp previously reported from a different *H. zea* strain (Perera et al. 2016; supplementary table S5). Size variation was accounted for by four indels. Specifically, our assembly has a predicted 5 bp insertion at positions 11,662 to 11,666 located adjacent to *trnS2* that resulted from an AACTA duplication that accounts for the discrepancy in position of this tRNA between the two assemblies. Compared to KJ930516.1, *rpnL* in our mitochondrial genome assembly has a 10 bp insertion at positions 12,777 to 12,886 comprising five AT repeat units, and an AATATT deletion from 13,544 to 13,548. This comparison to KJ930516.1 additionally predicts that our assembly has an AT dinucleotide deletion and nine substitutions within the AT-rich control region. The GC-content in both assemblies was about 19%, and typical of insect mitochondrial genomes (Crozier and Crozer 1999). Order and orientation of the 13 protein-coding, 22 tRNA, and 2 rRNA genes in our assembly (supplementary fig. S1 and table S5) were typical of lepidopteran mitochondrial genomes, and identical to the previous *H. zea* assembly (Perera et al. 2016). Specifically, *trnM* was inverted compared to the consensus animal mitochondrial genome (Boore 1999) and a non-canonical Arginine CGA codon putatively functions in initiation of cytochrome c oxidase I *(coxI)* translation. No variation was detected within tRNA or protein coding genes compared to KJ930516.1. Absence of mitochondrial-derived k-mers in the Illumina k-mer database after two rounds of polishing indicated high accuracy and confidence in variation between our mitochondrial assembly compared and the previous accession.

## Materials and Methods

### Contig assemblies from continuous long read (CLR) PacBio data

Adult *H. zea* were collected near the Mississippi State University Campus in Starkville, MS, USA during 2011 and maintained in a colony at the USDA-ARS Southern Insect Management Unit (SIMRU), Stoneville, MS, USA as described previously (Gore et al. 2005). Larvae were selected on a diagnostic dose of 2.0 μg ml^-1^ purified Cry1Ac, and survivors used to create the strain, HzStark_Cry1AcR. HzStark_Cry1AcRwas backcrossed every 5 generations to a susceptible line maintained at USDA-ARS SIMRU.

A single male pupa (homogametic with ZZ sex chromosome pair; ToLID ilHelZeax1) from HzStark_Cry1AcR was dissected laterally into eight ~20.0 μg sections. High molecular weight DNA was extracted from each section using the MagAttract HMW DNA Kit (Qiagen, Hilden, Germany) modified from manufacturer instructions to include gentle inversion to mix all components, an additional wash step, and eluting DNA from beads with three separate additions of 115.0 μl AE buffer at 37□C. Mean fragment size and quantity was estimated on a TapeStation 4200 (Agilent, Santa Clara, CA, USA; 100 bp to 48.5 kb ladder). HMW genomic DNA was sheared with a Covaris g-TUBE and a PacBio library was prepared using the SMRTbell Express Template Prep Kit 2.0 for Continuous Long Read (CLR) generation (Pacific Biosciences, Menlo Park, CA, USA). The library was size selected on a BluePippin (Sage Sciences, Beverly, MA, USA) with a 15.0 kb cutoff. The library was loaded on an 8M SMRTcell at a concentration of 35.0 pmol and run on a Sequel II (Pacific Biosciences) with Instrument Control Software Version 7.0 for a 20-hour movie time. Libraries were created from DNA of the same HzStark_Cry1AcR male using the TruSeq DNA PCR-Free Low Throughput Library Prep Kit with TruSeq DNA UD Indexes (Illumina, San Diego, CA, USA) to yield a standard paired end library with a 350±50 bp insert size. Paired end 150 bp sequence reads were generated on a NovaSeq 6000 (Illumina) at the Hudson Alpha Genome Sequencing Center (Huntsville, AL, USA). Genome size, repetitive content, and heterozygosity were estimated from 21-mer histograms of the paired-end Illumina reads generated with jellyfish (Marçais et al. 2011) and modeled in GenomeScope 2.0 (Ranallo-Benavidez et al. 2020).

For genome assembly, raw CLR subread BAM files were converted to FASTQ format using BamTools v2.5.1 (Barnett et al. 2011), then used as input for the FALCON assembler (Chin et al. 2016) using the pb-assembly conda environment v.0.0.8.1 (Pacific Biosciences; default parameters). FALCON-Unzip created primary and alternate contigs with one round of haplotype-aware polishing by Arrow (Pacific Biosciences). Duplicated sequence at the ends of primary contigs were removed with purge_dups v1.2.5 (Guan et al. 2020), with cutoffs estimated from an automatically generated histogram of k-mers. Purged primary sequences were added to the alternate haplotype contigs and purged of duplicates again.

### Mitochondrial genome assembly

The mitochondrial genome was assembled and polished using a custom workflow, mitoPolishCLR (Formenti and Stahlke 2022), modified from mitoVGP (Formenti et al. 2021a) and improved for arthropods. Raw CLR reads were filtered to remove those > 1.5-times the known *H. zea* mitochondrial genome size (15,343 bp; Perera et al. 2016; GenBank accession KJ930516.1), thereby reducing low quality reads. Length filtered reads were retained if they shared homology to KJ930516.1 within results from BLASTn searches (default parameters; Camacho et al. 2009), then assembled using Canu v.2.2 (Koren et al. 2017) with parameters adjusted for organelle assemblies. Contig polishing was performed with FreeBayes, then evaluated by QV scores generated using Merqury (Rhie et al. 2020) and a 31-mer database of Illumina short reads generated by Meryl (Miller et al. 2008). After a final round of trimming the mitochondrial assembly was linearized and annotated with MitoFinder (Allio et al. 2020). Start and stop codons for protein-coding genes were manually reviewed according to guidelines for mitochondrial genomes of lepidoptera (Cameron 2014).

### Scaffolding

Hi-C libaries were prepared at Dovetail Genomics (Santa Cruz, CA, USA) as prevously described (Lieberman-Aiden et al. 2009) from *H. zea* derived from the Benzon Reasearch colony (Carlisle, PA, USA) that were reared at the USDA-APHIS Forest Pest Methods Laboratory (Buzzards Bay, MA, USA). These libraries were sequenced at Dovetail on an Illumina HiSeq X (Illumina).

Hi-C data were then used for scaffolding by aligning them to contigs using bwa-mem v2.2.1 (Li 2013) with the −5SP options to allow chimeric read alignment. Matlock Hi-C processing (https://github.com/phasegenomics/matlock) and Juicebox v1.11.08 assembly tools (https://github.com/phasegenomics/juicebox_scripts) were used to convert the resulting BAM file and prepare the primary contig assembly for review. A manually curated contact-map relating the primary contigs to the aligned Hi-C reads was created in Juicebox v1.11.08. With the Hi-C contact map, we manually broke mis-joins, re-oriented inverted contigs, and joined contigs into scaffolds. The manually curated scaffold assembly was converted back to FASTA format for final polishing.

After scaffolding, the primary, alternate, and mitochondrial assemblies were merged and polished with one additional round of Arrow (Pacific Biosciences), followed by two rounds of variant identification using FreeBayes v1.0.2 (Garrison and Marth 2012). FreeBayes identified variants were filtered via Merfin (Formenti et al. 2021b) before being incorporated into a new polished consensus. The second round of Arrow-identified variants were also filtered via Merfin as the PacBio CLR and Illumina reads came from the same *H. zea* individual. Genome assembly quality scores were generated via Merqury (Rhie et al. 2020) before, between, and after each round of polishing to assess quality improvements. The polishCLR pipeline used here (https://github.com/isugifNF/polishCLR) is a reproducible Nextflow workflow (Di Tommaso et al. 2017) inspired by the Vertebrate Genome Project assembly pipeline (Rhie et al. 2021).

Contigs derived from non-*H. zea* biological contamination (such as microbial symbionts) were screened using the BlobToolkit (Challis et al. 2020) which implements BLAST+ (Camacho et al. 2009) and Diamond (Buchfink et al. 2015), and parsed using a python script (https://github.com/sheinasim/FindContaminantsFromBlob). Final assembly completeness was assessed by BUSCO v.5.2.2 with the lepidoptera_odb10 gene set of 5,286 orthologs (Simao et al. 2015; Seppey et al. 2019; Manni et al. 2021), and composition evaluated by a 21-mer k-mer spectra using KAT v.2.4.2 (Mapleson et al. 2016).

### Genome annotation

Repeats were identified using RepeatMasker v4.1.0 (Smit et al. 2005) using a combined repeat library built with *de novo* repeats using RepeatModeler v2.0.2 (Smit et al. 2008-2015) and lineage-specific Dfam v3.1 databases for Insecta and Lepidoptera (Storer et al. 2021). Scaffolded contigs from the primary assembly were submitted to the National Center for Biotechnology Information (NCBI) for automated eukaryotic genome annotation (Thibaud-Nissen et al. 2016).

## Supplementary Materials

Supplementary data are available from *Genome Biology and Evolution* online.

## Acknowledgements

This work was supported by the U.S. Department of Agriculture, Agricultural Research Service (USDA-ARS). The genome assembly was generated as part of the USDA-ARS Ag100Pest Initiative. The HzStark_Cry1AcR pupae for genome assembly were provided by USDA-ARS Southern Insect Management Research Unit (SIMRU). Funding for PacBio CLR read data was provided by the USDA-ARS Corn Insects & Crop Genetics Research Unit project number 5030-22000-019-00D. This research used resources provided by the SCINet project of the USDA-ARS project number 0500-00093-001-00-D. Jennifer Chang was supported, in part, by an appointment to the Research Participation Program at USDA-ARS, administered by the Oak Ridge Institute for Science and Education (ORISE) through an interagency agreement between the U.S. Department of Energy and USDA-ARS under contract number DE-AC05-06OR23100. Luke Tembrock was supported, in part, by a USDA-APHIS-PPQ cooperative agreement AP18PPQS&T00C074 to Colorado State University. Thanks members of the USDA-ARS Ag100Pest Team for sequencing and analysis support. Thanks to Sheron Simpson from the USDA-ARS Jamie Whitten Delta States Research Center, Genomics and Bioinformatics Research Unit for library preparation and sequencing of PacBio CLR libraries. Thanks to Calvin Pierce, USDA-ARS, SIMRU, Stoneville, MS for maintaining the HzStark_Cry1AcR colony, and Hannah Nadel and Lara Trozzo at the USDA Forest Pest Methods Laboratory in Buzzards Bay, MA for rearing the *H. zea* used to generate Hi-C libraries. All opinions expressed in this paper are the author’s and do not necessarily reflect the policies and views of USDA. Mention of trade names or commercial products in this publication is solely for the purpose of providing specific information and does not imply recommendation or endorsement by USDA. USDA is an equal opportunity provider and employer.

## Author Contributions

B.C., L.T., B.S., A.C., S.S., S.G., T.G., and K.H. designed and supervised the project; B.C., O.P., and L.T. prepared samples; A.S. J.C., S.C., and S.S analyzed data; B.C., A.S., and L.T. interpreted results; B.C., A.S., and L.T. wrote a majority of the manuscript; All authors read and approved the final manuscript.

## Data Availability

Raw WGS CLR and Illumina sequence data were deposited at DDBJ/ENA/GenBank within BioProject PRJNA804956, under the Sequence Read Archive (SRA) accessions SRR17965731 and SRR17993835, respectively. Raw Hi-C Illumina sequence data were deposited within BioProject PRJNA788876, under SRA accession SRR17229576. The annotated primary assembly version ilHeaZeax1.1 accession GCF_022581195.2 (BioProject PRJNA807638; Annotation Release 100) and ilHeaZeax1.1 alternate haplotype assembly versions accession GCA_022581175.1 (BioProject PRJNA807637) are available at NCBI. Both are under the Ag100Pest umbrella project, BioProject PRJNA555319. The primary assembly and annotations are also available at the i5k Workspace@NAL (Poelchau et al. 2015). The annotated mitochondrial genome assembly is available in the GenBank accession OM990843.1.

## Notes

https://www.ncbi.nlm.nih.gov/assembly/GCF_022581195.2/

### Competing Interest Statement

The authors have declared no competing interest.

