## supplementary for "A chromosome-scale genome assembly of a *Bacillus thuringiensis* Cry1Ac insecticidal protein resistant strain of *Helicoverpa zea*"

Supplementary data

**Table S1**

Data used to generate the *Helicoverpa zea* ilHelZeax1.1 genome assembly.

|  |  |  | Genomic reads | | | |  |  |  |
| --- | --- | --- | --- | --- | --- | --- | --- | --- | --- |
| Library name | Source |  | Platform | Type | Count (M) | Gb |  | Use^3^ | SRA accession^4^ |
| CEWMale14 | Pupa ♂14^1^ |  | Pacific Biosystems Sequel I | CLR | 5.7 | 54.0 |  | C, P | SRR17965731 |
| CEWmale14_Illumina | Pupa ♂14^1^ |  | Illumina NovaSeq 6000 | 2x150 bp | 154.6 | 23.3 |  | P | SRR17993835 |
| DTG_HiC_967 | Pupa 2nd^2^ |  | Illumina HiSeq X | 2x150 bp | 71.2 | 21.3 |  | S | SRR17229576 |

1. Bt Cry1Ac resistance colony (Cry1AcR) maintained at USDA-ARS, Southern Insects Management Research Unit, Stoneville, MS.
2. Sex not determined; individual from a commercial colony sold by Benson Research Inc. (Carlisle, PA, USA).
3. Use code: contig assembly (C), contig polishing (P), and scaffolding (S).
4. Under the USDA-ARS Ag100Pest umbrella project NCBI accession [PRJNA555319](https://www.ncbi.nlm.nih.gov/bioproject/555319).

**Table S2**

Genome assembly versions generated for the *Helicoverpa zea* strain HzStark_Cry1AcR compared to previous assembles from the genus *Helicoverpa*. There were no gaps (Ns) in contig assemblies. Benchmarking Universal Single-Copy Orthologs (BUSCOs) from the lepidoptera_odb10 orthologs (*n* = 5286) with NA (not applicable) indicated for alternate assemblies.

|  |  |  | |  |  | | ilHelZeax1.1 | |  | Short read assemblies | |  | Hybrid assembly |
| --- | --- | --- | --- | --- | --- | --- | --- | --- | --- | --- | --- | --- | --- |
|  |  | Initial contigs | |  | De-duplicated contigs | |  | Scaffolds |  | Hzea_1.0^3^ | Harm_1.0^4^ |  | ASM1716586v1^5^ |
| Contig/Scaffold | | Primary | Alternate |  | Primary | Alternate^1^ |  | Primary^2^ |  | Scaffolds | Scaffolds |  | Scaffolds |
|  | Number | 134 | 765 |  | 81 | 801 |  | 42 |  | 2,975 | 998 |  | 1,211 |
|  | Length (bp) | 412,401,926 | 99.4 |  | 375,173,893 | 133,873,018 |  | 375,165,593 |  | 341,147,348 | 337,087,553 |  | 390,016,491 |
|  | GC content (%) |  |  |  |  | 36.89 |  | 36.86 |  | 32.50 | 32.15 |  | 36.55 |
|  | Gaps | 0 | 0 |  | 0 | 0 |  | 49 |  | 32,923 | 26,906 |  | 0 |
|  | N bases | 0 | 0 |  | 0 | 0 |  | 4900 |  | 34,741,529 | 37,110,157 |  | 0 |
|  | N50 (bp) | 8.2,8173 | 0.343 |  | 8.5 | 464,725 |  | 12,841,304 |  | 201,477 | 1,000,414 |  | 456,792 |
|  | L50 | 18 | 87 |  | 17 | 90 |  | 14 |  | 469 | 93 |  | 276 |
|  | N90 (bp) | 1,919,501 | 0.416 |  | 2.8 | 49,392 |  | 9,278,413 |  | 52,272 | 175,335 |  | 177,008 |
|  | L90 | 56 | 449 |  | 44 | 417 |  | 27 |  | 1,755 | 417 |  | 805 |
|  | Max (Mb) | 15,903,638 |  |  |  | 1,812,199 |  | 18,805,280 |  | 1,847,547 | 6,146,627 |  | 1,683,735 |
|  | BUSCOs |  |  |  |  |  |  |  |  |  |  |  |  |
|  | Complete (C) | 5230 (99.0%) | NA |  | 5226 (98.8%) | NA |  | 5227 (98.8%) |  | 5110 (96.6%) | 5216 (98.7%) |  | 5084 (96.1%) |
|  | Single copy (S) | 4713 (89.2%) | NA |  | 5209 (98.5%) | NA |  | 5209 (98.5%) |  | 5045 (95.4%) | 5185 (98.1%) |  | 4500 (85.1%) |
|  | Duplicated (D) | 517 ( 9.8%) | NA |  | 18 ( 0.3%) | NA |  | 18 ( 0.3%) |  | 65 ( 1.2%) | 31 ( 0.6%) |  | 584 (11.0%) |
|  | Fragmented (F) | 17 ( 0.3%) | NA |  | 17 ( 0.3%) | NA |  | 17 ( 0.3%) |  | 56 ( 1.1%) | 21 ( 0.4%) |  | 76 ( 1.4%) |
|  | Missing (M) | 39 ( 0.7%) | NA |  | 42 ( 0.9%) | NA |  | 42 ( 0.9%) |  | 120 ( 2.3%) | 49 ( 0.9%) |  | 126 2.5%) |

1. WGS Project: JAKYJX01; GenBank accession: GCA_022581175.1 *(this assembly)*
2. WGS Project: JAKYJW01; GenBank accession: GCF_022581195.2 *(this assembly)*
3. WGS Project: NFMG01; GenBank accession: GCA_002150865.1; Pierce et al. (2017)
4. WGS Project: NHMN01; GenBank accession: GCA_002156985.1; Pierce et al. (2017)
5. WGS Project: JACGXX01; GenBank accession: GCA_017165865.1; National Institute of Crop Science, RDA (unpublished)

**Table S3**

Comparison of the *Helicoverpa zea* strain HzStark_Cry1AcR scaffolded assembly to other chromosome-level assemblies from species in the lepidopteran Family Noctuidae. NA = not available.

| Metric | | ilHelzeax1.1^1^ | MamBras1.0^2^ | CraLigu1.1^3^ | ZJU_Sfrug_1.0^4^ | NJAU_Sex_v1^5^ | AutGamm1.1^6^ |
| --- | --- | --- | --- | --- | --- | --- | --- |
| Contig assembly | |  |  |  |  |  |  |
|  | Total number | 65 | 70 | 95 | 617 | 667 | 130 |
|  | Length (bp) | 375,163,385 | 576,175,243 | 437,993,611 | 486,287,157 | 446,801,326 | 373,067,087 |
|  | N50 (Mb) | 10,841,041 | 17,758,554 | 12,212,030 | 1,129,999 | 3,467,000 | 11,185,469 |
|  | L50 | 17 | 14 | 15 | 129 | 31 | 15 |
|  | N90 (bp) | 3,915,976 | 7,694,024 | 2,987,509 | 380,677 | 514,962 | 3,829,268 |
|  | L90 | 34 | 33 | 43 | 423 | 159 | 33 |
|  | Largest contig (bp) | 15,512,169 | 23,412,814 | 18,426,615 | 22,364,714 | 15,878,684 | 19,124,284 |
| Primary scaffold assembly | |  |  |  |  |  |  |
|  | Total number | 42 | 43 | 33 | 92 | 301 | 103 |
|  | Length (bp) | 375,165,593 | 576,175,243 | 437,993,611 | 486,287,157 | 446,801,326 | 373,067,087 |
|  | Gaps | 49 | 27 | 62 | 525 | 370 | 27 |
|  | GC content (%) | 36.86 | 38.31 | 36.03 | 36.43 | 36.64 | 35.66 |
|  | N bases | 4900 | 8900 | 17800 | 52500 | 335,524 | 7600 |
|  | N50 (bp) | 12,841,304 | 19,384,296 | 15,030,495 | 16,346,893 | 14,363,500 | 12,167,202 |
|  | L50 | 14 | 14 | 14 | 14 | 15 | 14 |
|  | N90 (bp) | 9,278,413 | 13,925,230 | 10,234,456 | 10,588,980 | 8,256,880 | 6,891,251 |
|  | L90 | 27 | 26 | 27 | 28 | 29 | 29 |
|  | Largest contig (bp) | 18,805,280 | 35,467,487 | 21,317,134 | 16,139,548 | 19,740,469 | 15,294,816 |
|  | Chromosomes |  |  |  |  |  |  |
|  | Autosomes | 30 | 30 | 30 | 30 | 30 | 30 |
|  | Sex chromosome(s) | Z | Z | Z+W | Z+W | Z+W | Z+W |
|  | BUSCOs |  |  |  |  |  |  |
|  | Complete (C) | 5227 (98.8%) | 5230 (98.9%) | 5229 (98.4%) | 5057 (95.7%) | 5189 (98.1%) | 5224 (98.9%) |
|  | Complete single copy (S) | 5209 (98.5%) | 5203 (98.4%) | 5203 (98.4%) | 4265 (80.7%) | 5076 (96.0%) | 5210 (98.6%) |
|  | Complete duplicated (D) | 18 ( 0.3%) | 27 ( 0.5%) | 26 ( 0.5%) | 792 (15.0%) | 113 ( 2.1%) | 14 ( 0.3%) |
|  | Fragmented (F) | 17 ( 0.3%) | 13 ( 0.2%) | 14 ( 0.3%) | 21 ( 0.4%) | 17 ( 0.3%) | 13 ( 0.2%) |
|  | Missing (M) | 42 ( 0.9%) | 43 ( 0.9%) | 43 ( 0.8%) | 208 ( 3.9%) | 80 ( 1.6%) | 49 ( 0.9%) |
| Alternate assembly (pseudohaplotype) | |  |  |  |  |  |  |
|  | Total number | 801 | 292 | 1,204 | NA | NA | 183 |
|  | Length (bp) | 133,873,018 | 23,004,467 | 399,761,949 | NA | NA | 337,172,335 |
|  | N50 (bp) | 464,725 | 552,583 | 601,847 | NA | NA | 9,142,168 |
|  | L50 | 90 | 14 | 204 | NA | NA | 15 |
|  | N90 (bp) | 49,392 | 26215 | 175436 | NA | NA | 2443531 |
|  | L90 | 417 | 182 | 659 | NA | NA | 38 |
|  | Largest contig (bp) | 1,812,199 | 1,612,378 | 6,028,452 | NA | NA | 13,444,919 |

1. *Helicoverpa zea* WGS Project: JAKYJW01; GenBank accession: GCF_022581195.2 *(this assembly)*
2. *Mamestra brassicae*; WGS Project: CAJHZQ01; GenBank accession: GCA_905163435.1 (Welcome Sanger Institute, unpublished)
3. *Craniophora ligusti*; GenBank accession: GCA_905163465.1 (Welcome Sanger Institute, unpublished)
4. *Spodoptera frugiperda*; WGS Project: WMCG01; GenBank accession: GCA_011064685.1; Xiao H et al. 2020
5. *Spodoptera exigua*; WGS Project: WNNL01; GenBank accession: GCA_011316535.1 ; Zhang et al. 2020
6. *Autographa gamma*; WGS Project: CAJHUD01; GenBank accession: GCA_905146925.1 (Welcome Sanger Institute, unpublished)

**Table S4**

*Helicoverpa zea* genome assembly and annotation metrics; A total of 110,825,260 bp (29.5%) of the assembly was masked.

| Repeat type | | Count | Length (bp) | Proportion |
| --- | --- | --- | --- | --- |
| *Retroelements (SINE + P + LINE + LTR)* | | *173,402* | *42,171,018* | *11.24 %* |
|  | SINEs: | 63,185 | 12,139,546 | 3.24 % |
|  | Penelope (P) | 1,837 | 301,907 | 0.08 % |
|  | LINEs: | 96,647 | 24,042,822 | 6.41 % |
|  | CRE/SLACS | 0 | 0 | 0.00 % |
|  | L2/CR1/Rex | 18,385 | 5,238,677 | 1.40 % |
|  | R1/LOA/Jockey | 18,409 | 6,526,900 | 1.74 % |
|  | R2/R4/NeSL | 1,552 | 834,716 | 0.22 % |
|  | RTE/Bov-B | 41,448 | 7,290,424 | 1.94 % |
|  | L1/CIN4 | 47 | 15,252 | 0.01 % |
|  | Others | 16,806 | 4,136,853 | 1.1 % |
|  | Long terminal repeat (LTR) elements | 11,733 | 5,687,324 | 1.54 % |
|  | BEL/Pao | 1,589 | 893,460 | 0.24 % |
|  | Ty1/Copia | 863 | 850,644 | 0.23 % |
|  | Gypsy/DIRS1 | 8,175 | 3,779,507 | 1.01 % |
|  | Retroviral | 0 | 0 | 0.00 % |
|  | Others | 1,106 | 165,713 | 0.04 % |
| *DNA Transposons* | | *40,016* | *10,750,078* | *2.87 %* |
|  | hobo-Activator | 10,242 | 3,020,129 | 0.80 % |
|  | Tc1-IS630-Pogo | 10,056 | 3,077,115 | 0.82 % |
|  | En-Spm | 0 | 0 | 0.00 % |
|  | MuDR-IS905 | 0 | 0 | 0.00 % |
|  | PiggyBac | 4,251 | 1,867,284 | 0.50 % |
|  | Tourist/Harbinger | 1,978 | 461,113 | 0.12 % |
|  | Mirage, P-element, Transib | 514 | 214,464 | 0.06 % |
|  | Other | 12,975 | 2,109,973 | 0.57 % |
| Helitron (Rolling circle) | | 128,948 | 17,517,321 | 4.67 % |
| Unclassified | | 156,092 | 35,918,128 | 9.57 % |
| Total interspersed repeats | | 498,458 | 88,537,898 | 23.60 % |
| Small RNAs | | 14,221 | 1,988,953 | 0.53 % |
| Satellites: | | 3267 | 223,839 | 0.06 % |
| Simple repeats | | 89,467 | 3,849,438 | 1.03 % |
| Low complexity | | 14,690 | 675,123 | 0.18 % |

**Table S5**

Gene coding regions of the *Helicoverpa zea* mitochondrial genome.

| Accession KJ930516.1 (Perera et al. 2016) | | | | | | | Accession OM990843.1 *(This assembly)* | | | | | | | | | |
| --- | --- | --- | --- | --- | --- | --- | --- | --- | --- | --- | --- | --- | --- | --- | --- | --- |
| **Gene** | | | | **Codons** | | **tRNA** | **Gene** | | | | **Codons** | | | | | **tRNA** |
| **Name** | **Start** | **Stop** | **Len** | **Init Term** | | **anticodon** | **Start Stop** | | **Len** | | | **Init Term** | | | **Anticodon** | |
| *trnM* | 1 | 69 | 69 |  |  | CAT | 1 | 69 |  | | |  | |  | CAT | |
| *trnI* | 70 | 133 | 64 |  |  | GAT | 70 | 133 |  | | |  | |  | GAT | |
| *trnQ* | 199 | 131 | 69 |  |  | TTG | 199 | 131 |  | | |  | |  | TTG | |
| *nad2* | 245 | 1,255 | 1,011 | ATT | TAA |  | 245 | 1,255 |  | | | ATT | | TAA |  | |
| *trnW* | 1,258 | 1,325 | 68 |  |  | TCA | 1,258 | 1,325 |  | | |  | |  | TCA | |
| *trnC* | 1,386 | 1,318 | 69 |  |  | GCA | 1,386 | 1,318 |  | | |  | |  | GCA | |
| *trnY* | 1,462 | 1,397 | 66 |  |  | GTA | 1,397 | 1,462 |  | | |  | |  | GTA | |
| *cox1* | 1,465 | 2,995 | 1,531 | CGA | T+polyA |  | 2,995 | 1,465 |  | | | CGA | | T+polyA |  | |
| *trnL2* | 2,996 | 3,062 | 67 |  |  | TAA | 2,996 | 3,062 |  | | |  | |  | TAA | |
| *cox2* | 3,063 | 3,744 | 682 | ATG | T+PolyA |  | 3,063 | 3,744 |  | | | ATG | | T+polyA |  | |
| *trnK* | 3,745 | 3,815 | 71 |  |  | CTT | 3,745 | 3,815 |  | | |  | |  | CTT | |
| *trnD* | 3,820 | 3,885 | 66 |  |  | GTC | 3,820 | 3,885 |  | | |  | |  | GTC | |
| *atp8* | 3,886 | 4,047 | 162 | ATC | TAA |  | 3,886 | 4,047 |  | | | ATC | | TAA |  | |
| *atp6* | 4,041 | 4,718 | 678 | ATG | TAA |  | 4,041 | 4,718 |  | | | ATG | | TAA |  | |
| *cox3* | 4,718 | 5,503 | 786 | ATG | TAA |  | 4,718 | 5,503 |  | | | ATG | | TAA |  | |
| *trnG* | 5,506 | 5,571 | 66 |  |  | TCC | 5,506 | 5,571 |  | | |  | |  | TCC | |
| *nad3* | 5,572 | 5,925 | 354 | ATT | T+polyA |  | 5,572 | 5,923 |  | | | ATT | | T+polyA |  | |
| *trnA* | 5,924 | 5,992 | 69 |  |  | TGC | 5,924 | 5,992 |  | | |  | |  | TGC | |
| *trnR* | 5,993 | 6,059 | 67 |  |  | TCG | 5,993 | 6,059 |  | | |  | |  | TCG | |
| *trnN* | 6,087 | 6,152 | 66 |  |  | GTT | 6,087 | 6,152 |  | | |  | |  | GTT | |
| *trnS1* | 6,154 | 6,221 | 68 |  |  | GCT | 6,155 | 6,220 |  | | |  | |  | GCT | |
| *trnE* | 6,222 | 6,288 | 67 |  |  | TTC | 6,222 | 6,288 |  | | |  | |  | TTC | |
| *trnF* | 6,353 | 6,287 | 67 |  |  | GAA | 6.353 | 6.287 |  | | |  | |  | GAA | |
| *nad5* | 8,099 | 6,354 | 1,746 | ATT | TAA |  | 8,099 | 6,354 |  | | | ATT | | TAA |  | |
| *trnH* | 8,166 | 8,100 | 67 |  |  | GTG | 8,166 | 8,100 |  | | |  | |  | GTG | |
| *nad4* | 9,505 | 8,167 | 1,339 | ATG | T+PolyA |  | 9,505 | 8,167 |  | | | ATG | | T+PolyA |  | |
| *nad4l* | 9,840 | 9,550 | 291 | ATG | TAG |  | 9,843 | 9,550 |  | | | ATA | | TAG |  | |
| *trnT* | 9,844 | 9,909 | 66 |  |  | TGT | 9,844 | 9,909 |  | | |  | |  | TGT | |
| *trnP* | 9,974 | 9,910 | 65 |  |  | TGG | 9,974 | 9,910 |  | | |  | |  | TGG | |
| *nad6* | 9,982 | 10,515 | 534 | ATT | TAA |  | 9,982 | 10,515 |  | | | ATT | | TAA |  | |
| *cob* | 10,516 | 11,667 | 1,152 | ATG | TAA |  | 10,516 | 11,667 |  | | | ATG | | TAA |  | |
| *trnS2* | 11,673 | 11,738 | 66 |  |  | TGA | 11,678 | 11,743 |  | | |  | |  | TGA | |
| *nad1* | 12,700 | 11,762 | 939 | ATG | TAA |  | 12,700 | 11,762 |  | | | ATG | | TAA |  | |
| *trnL1* | 12,769 | 12,702 | 68 |  |  | TAG | 12,769 | 12,702 |  | | |  | |  | TAG | |
| *rrnL* | 14,159 | 12,770 | 1,390 |  |  |  | 14,164 | 12,770 |  | | |  | |  |  | |
| *trnV* | 14,225 | 14,160 | 66 |  |  | TAC | 14,230 | 14,165 |  | | |  | |  | TAC | |
| *rrnS* | 15,019 | 14,226 | 794 |  |  |  | 15,024 | 14,231 |  | | |  | |  |  | |
| Control | 15,020 | 15,348 | 329 |  |  |  | 15,025 | 15,351 | |  |  | |  | | |  |
| *trnI** | 15,173 | 15,268 |  |  |  | AAT |  |  | |  |  | |  | | |  |

**Fig. S1** Annotation of contig-based assembly of the 15,351 bp *Helicoverpa zea* mitochondrial genome. Arrows indicate order and orientation of genic regions (green), with major (+) and minor (-) strands indicated by clockwise and counterclockwise arrows, respectively. Coding regions for proteins (yellow), ribosomal RNAs (red) and tRNA (pink) correspondingly indicated. The AT-rich origin of replication is shown in blue.

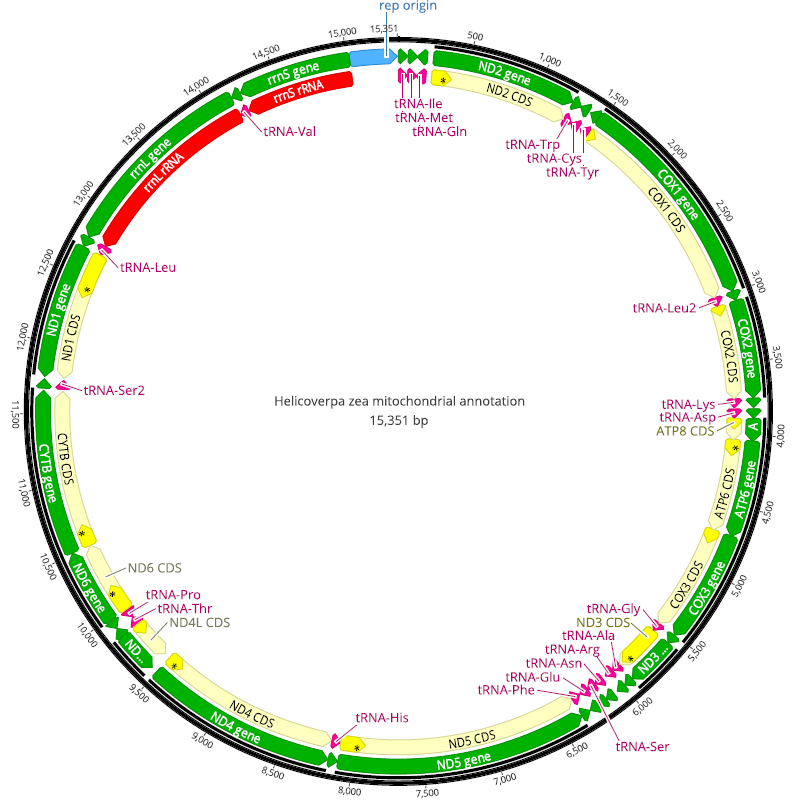
